# *DNM2*-CMT neuropathy stems from disrupted Schwann cell function and shows limited therapeutic reversibility

**DOI:** 10.64898/2025.12.02.691765

**Authors:** Marie Goret, Thomas Arbogast, Jocelyn Laporte

## Abstract

Dominant loss-of-function mutations in *DNM2* cause Charcot-Marie-Tooth (CMT) neuropathy characterized by sensory and motor deficits associated with myelin and/or axonal abnormalities and muscle atrophy. Increasing DNM2 activity from embryogenesis has been reported to ameliorate neuromuscular phenotypes in the *Dnm2^K562E/+^* CMT mouse; however, this model displays predominantly muscle pathology and limited nerve involvement, precluding rigorous evaluation of neuropathic mechanisms and potential therapies. Here, we performed comprehensive behavioral, electrophysiological, histological and molecular analyses to characterize the *Dnm2^K562E/^*^SC*-*^ mouse, which combines systemic heterozygosity for the common K562E mutation together with Schwann cell (SC)-specific deletion of wild-type *Dnm2*. This model faithfully reproduces key clinical and pathological features of *DNM2*-CMT, including motor deficits, reduced general force and coordination, and severe sensory and motor conduction deficits associated with axonal loss, demyelination, and inflammation. Mechanistically, we delineate a coherent pathological sequence that explains the profound functional deficits. In particular, a downregulation of the transcription factor EGR2, a master regulator of myelin gene expression, and of the myelin protein MPZ correlates with demyelination. To evaluate the therapeutic potential of DNM2 supplementation, post-symptomatic intrathecal delivery of AAV9-DNM2 driven by the Schwann cell-specific MPZ promoter was performed at 4 weeks. Although DNM2 expression increased in peripheral nerves (∼1.9-fold), no significant improvements were observed across behavioural, electrophysiological, structural, or molecular parameters. Together, these findings establish the *Dnm2^K562E/^*^SC*-*^ mouse as a robust preclinical model, recapitulating key features of *DNM2-*CMT, and provides crucial insight into the biological and temporal constraints that must guide future therapeutic strategies for *DNM2*-CMT.

## Introduction

Charcot-Marie-Tooth disease (CMT) is the most common inherited peripheral neuropathy, with more than 100 causative genes identified across demyelinating and axonal forms of the disorder^1^. Among these, *DNM2* encodes dynamin 2, a large GTPase involved in membrane trafficking and cytoskeletal dynamics^2–4^. Dominant *DNM2* mutations cause both intermediate (CMT-DIB; MIM#606482)^5,6^ and axonal (CMT2M)^7^ forms of CMT. Patients with *DNM2*-related CMT typically present with slowed nerve conduction velocities, impaired proprioception, decreased reflexes, progressive distal weakness and atrophy, gait disturbances, and skeletal deformities^8^. Additional features such as ptosis, ophthalmoparesis, cataracts, or neutropenia may occur^9–11^. Nerve biopsies commonly reveal loss of large myelinated fibers, clusters of regenerating axons, focal myelin thickening, increased g-ratio, occasional onion bulb formation and few fibers with Schmidt-Lanterman incisures^10–13^. Currently, no therapies are available^14,15^, underscoring the need for mechanistic and therapeutic studies.

The mechanisms underlying *DNM2*-related CMT are not fully understood. Several studies highlighted that *DNM2*-CMT most probably arise from a DNM2 dominant-negative loss-of-function mechanism. Mutations lead to decreased GTPase activity^16,17^, and complete *Dnm2* deletion in Schwann cells (SCs) resulted in severely impaired axonal sorting and myelination onset leading to a rapidly-developing peripheral neuropathy^18^. However, the *Dnm2^K562E/+^* mouse, carrying the most frequent patient mutation^19^, exhibit motor and muscle abnormalities but only subtle nerve pathology^20–22^. Consequently, this model offers limited insight into Schwann cell-specific mechanisms and hinders the evaluation of nerve-targeted therapeutic strategies. Previous preclinical studies have suggested that enhancing DNM2 activity could compensate for the loss-of-function mechanism underlying CMT. Indeed, genetic crosses that increased DNM2 activity from embryogenesis, either by introducing the hyperactive DNM2-S619L variant or reducing BIN1, a negative regulator of DNM2, rescued neuromuscular phenotypes in *Dnm2^K562E/+^* mice ^20,22^. However, these proof-of-concept strategies relied on genetic crosses that increased DNM2 activity ubiquitously from embryogenesis, limiting their translational potential. The feasibility of a postnatal, gene-supplementation-based approach to restore DNM2 function in peripheral nerves has not yet been explored.

To address these gaps, we performed here a detailed phenotypic characterization of the *Dnm2^K562E^*^/SC-^ mouse model, which combines systemic expression of the K562E mutant allele with Schwann cell (SC)-specific deletion of WT *Dnm2*^21^. This model exhibits a severe demyelinating neuropathy and provides a platform to dissect Schwann cell-autonomous disease mechanisms. We further evaluated whether targeted DNM2 augmentation after symptom onset could reverse neuropathic pathology by delivering AAV9-DNM2 under the Schwann cell-specific MPZ promoter via intrathecal injection, followed by comprehensive behavioral, electrophysiological, histological, and molecular assessments. *Dnm2^K562E/^*^SC*-*^ mice exhibited tremor, severe coordination deficits, conduction impairments, axonal loss, and demyelination associated with reduced MPZ and EGR2 expression and increased inflammation, thereby recapitulating a demyelinating neuropathy closely resembling that observed in patients. Importantly, DNM2 supplementation at 4 weeks of age failed to reverse these defects, indicating that DNM2 activity is required during early postnatal Schwann cell maturation and that late Schwann cell-specific DNM2 augmentation is insufficient to repair established pathology.

## Materials and methods

### Mouse models

*Dnm2^K562E/^*^SC*-*^ mice were generated through a two-step breeding strategy. First, MPZ-Cre⁺ mice (strain #017927, RRID:IMSR_JAX:017927; obtained from Charles River) were crossed with *Dnm2* exon 8 floxed mice (ICS project K546/IR2694)^23^. The resulting MPZ-Cre⁺ *Dnm2*^fl/+^ progeny (*Dnm2^+/^*^SC*-*^) were then crossed with *Dnm2^K562E/+^* mice^20,21^ to produce experimental animals. All mice, including controls, expressed MPZ-Cre, had a mixed genetic background (75% C57BL/6J and 25% C57BL/6N), included both sexes, and were analyzed at 8 weeks of age. Of note, MPZ-Cre expression initiates between embryonic days 13.5 and 14^24^.

For genotyping, primers “6115 Er KE” and “6116 Ef KE” were used to detect K562E mutation, primers “Cre 160” and “Cre 161” to detect the Cre, and “4613” and “4614” for the *Dnm2* exon 8 floxed allele (Supplementary Table 1 Reagents).

Mice were housed in ventilated cages, with unrestricted access to food and water. Environmental parameters were maintained at 19-22 °C and 40-60% relative humidity, under a 12-hour light/dark cycle. Breeding mice received SAFE® D03 diet, transitioning to SAFE® D04 post-weaning, while *Dnm2^K562E/^*^SC*-*^ mice remained on SAFE® D03 supplemented with DietGel® 76A (SKU: 72-07-5022) from 3 weeks.

### AAV vector design and production

Recombinant AAV9 vectors were produced by the Molecular Biology and Virus Facility at IGBMC using a standard triple transfection protocol in HEK293T/17 cells. Expression plasmid encoding murine *Dnm2* (transcript variant 3, NCBI RefSeq NM_007871.2) cDNA (excluding exons 12b and 13ter) was cloned under the control of rat MPZ promoter. Constructs (pAAV-rMPZ-mDNM2, pAAV-CMV-MCS) were co-transfected with the pHelper plasmid (Agilent, CA, USA) and the pAAV2/9 capsid plasmid (P0008, Penn Vector Core, PA, USA). Viral particles were harvested 48 hours post-transfection from Benzonase-treated (100 U/mL, Merck) cell lysates. Purification was performed via iodixanol gradient ultracentrifugation (OptiPrep™, Axis Shield), followed by dialysis and concentration in Dulbecco’s PBS supplemented with 0.5 mM MgCl₂, using Amicon Ultra-15 centrifugal filters (100 kDa cutoff, Merck Millipore). Final titers were determined by qPCR using LightCycler 480 SYBR Green I Master mix (Roche), with primers specific to mDNM2, or the CMV enhancer (Supplementary Table 1 Reagents). Purified vectors were aliquoted and stored at −80 °C until use.

### In vivo AAV injections

Intrathecal injections were carried out at 4 weeks of age under 1.5–2% isoflurane anesthesia. After shaving and disinfecting the lumbar region, 10 µL of AAV9 was injected into the subarachnoid space between lumbar vertebrae L5 and L6 using a 27G needle connected to a Hamilton syringe, to a dose of 3.1 × 10¹¹ gc per mouse. Correct needle placement was confirmed by tail-flick reflex. For untreated controls, equivalent volumes and concentrations of AAV-empty vectors were administered using the same injection protocol.

### Behavioral tests

Behavioral assessments were conducted at 7 weeks of age, on the same weekday and in the same order for all animals, by a single experimenter blinded to genotype and treatment.

The hanging test measured latency to fall from an inverted grid (max 60 s); three trials were performed per animal, and the two best were averaged. Gait analysis was performed on a motorized treadmill equipped with a camera, at a constant speed of 10cm/s. Body stretch (nose to tail base) and stride length (step distance normalized to body stretch) were quantified, three measures were averaged. Sensorimotor coordination was assessed using the notched bar test, where mice crossed a horizontal bar with alternating notches. The number of falls and hindlimb slips into notches were recorded over 10 trials per mouse and averaged.

Mechanical sensitivity was measured using the Von Frey test. Calibrated filaments (0.6 g to 4 g) were applied perpendicularly to the plantar surface of the hind paw, recording rapid foot lifting as a positive reaction and the 50% paw withdrawal threshold was determined using the up-down method^25^ and the up-down reader program^26^. Thermal sensitivity was measured using the hot plate (52 °C), recording latency to hind paw withdrawal or licking. For tail immersion (48 °C), latency to tail flick was measured. Mice were habituated to the restraint tube on day 1, and measurements were taken on day 2. Each test consisted of three trials with 5-minute intervals, and values were averaged. Body length was measured post-mortem (nose to tail base).

### Electrophysiology

Mice were anesthetized with isoflurane (induction at 3%, maintenance at 2%) in oxygen-enriched air. Body temperature was monitored via rectal probe and maintained between 34–37 °C using a heating pad and a heat lamp. Eyes were protected with Ocrygel to prevent drying. EMG was performed using a Natus Mag2Health system. For sensitive evaluation, stimulating and recording electrodes were placed on the tail (separated by 2cm), the ground electrode in between (Fig. 2). For motor nerve evaluation, needle electrodes were placed for proximal stimulation at the sciatic nerve and distal stimulation at the ankle. The ground electrode was inserted subcutaneously on the contralateral side, and recordings were obtained from the plantar muscle (Fig. 2). Stimulation duration was 0.1 ms, and intensity was adjusted to elicit maximal responses.

### *In situ* muscle force

Contractile properties of the TA muscle were assessed in situ at 8 weeks of age using the Aurora Scientific 1300A system. Anesthesia was induced by intraperitoneal injection of domitor (2 mg/kg), fentanyl (0.28 mg/kg), and diazepam (8 mg/kg). Mice were placed under a heat source to maintain body temperature. The TA muscle was surgically exposed, and the distal tendon was attached to an isometric force transducer. The sciatic nerve was accessed through a small incision at the thigh, and stimulating electrodes were placed either around the nerve or directly on the muscle. Muscle length and stimulation intensity were optimized for maximal twitch force. Isometric tetanic force was first recorded following sciatic nerve stimulation (150 Hz, 0.5 s duration), followed by force-frequency measurements (1–150 Hz). These tests were repeated with direct muscle stimulation and a fatigue protocol was conducted using 80 consecutive stimulations (40 Hz, 1 s on / 3 s off). Peak force was recorded for each contraction. For fatigue analysis, tetanic force was averaged every five contractions, and fatigue was expressed as the percent drop in force from the first five to the last five contractions. Specific force (mN/mg) was calculated by normalizing force to TA muscle mass.

### Tissue collection

Muscles were dissected and weighed. TA muscles were snap-frozen in liquid nitrogen-cooled isopentane and stored at −80 °C for histological or protein analyses. Sciatic nerves for molecular studies were snap-frozen in liquid nitrogen and stored at −80 °C. For fiber bundle isolation (bungarotoxin immunofluorescence), the entire hindlimb (cut above the knee) was fixed with the leg extended in 4% PFA for 24 h at 4 °C. The extensor digitorum longus (EDL) muscle was then dissected, rinsed in PBS (3x), incubated in 25% sucrose at 4 °C until the tissue sank, and stored at −80 °C. Sciatic nerves for semi-thin sections were fixed in 2.5% glutaraldehyde and 2.5% paraformaldehyde in 0.1 M cacodylate buffer (pH 7.4) and stored at 4 °C. For immunofluorescence, nerves were fixed flat on filter paper in 4% PFA for 24 h at 4 °C, rinsed in PBS (3×), incubated in 25% sucrose at 4 °C until the tissue sank, and stored at −80 °C.

### Muscle histology

8µm transversal TA sections were cut using a cryostat and stained with HE by the IGBMC histology platform. Slides were scanned using the Carl Zeiss Axioscan 7, and 20X images were used for analysis. Fiber segmentation was performed on HE stained sections using CellPose^27^ software. MinFeret diameters were quantified in Fiji^28^. One transversal section of the whole muscle was quantified for each animal.

### Sciatic nerve histology

Fixed sciatic nerves samples were washed in cacodylate buffer (0.1 M, pH 7.4) for 30 minutes. Post-fixation was performed with 1% osmium tetroxide in 0.1 M cacodylate buffer for 1 hour at 4 °C. Samples were subsequently dehydrated through an ascending ethanol series (50%, 70%, 90%, 100%) and treated with propylene oxide, 30 minutes per step. Nerves were oriented transversely and embedded in Epon 812. Semithin sections (2 µm thick), performed as distally as possible, were stained with toluidine blue. Images were acquired at 63X magnification using a Zeiss Axio Observer 7 microscope. Axon categories and myelin abnormalities were classified across the entire nerve section using QuPath^29^. g-ratio, axon diameter, and myelin thickness were quantified on the whole stained section using the AxonDeepSeg segmentation software^30^.

### Muscle immunofluorescence

For bungarotoxin staining, EDL muscles were thawed on ice, and single fibers or small bundles were manually isolated under a stereomicroscope in cold PBS. Samples were mounted on slides, fixed in 4% PFA for 20 minutes, then permeabilized in PBS containing 0.5% Triton X-100 for 30 minutes at RT. Blocking was performed for 1 hour in PBS with 1% BSA and 0.1% Triton X-100. Slides were incubated with bungarotoxin antibody and DAPI (Supplementary Table 1 Reagents) for 1.5 hours at RT, then mounted with ProLong Gold Antifade. Images were acquired at 20X using a Zeiss Axio Observer 7 microscope. Neuromuscular junctions were detected on Fiji by adjusting the threshold and signal area was measured.

### Sciatic nerve immunofluorescence

Longitudinal 10 µm sciatic nerve sections were fixed in 4% paraformaldehyde for 20 minutes, permeabilized with PBS containing 0.2% Triton X-100 (Merck #T8787-250ML) for 10 minutes, and blocked for 1 hour in PBS with 0.1% Triton X-100 and 5% BSA (MP Bio #02160069-CF). Sections were incubated overnight at 4 °C with primary antibodies (Supplementary Table 1 Reagents). Alexa Fluor-conjugated secondary antibodies, along with DAPI, were applied for 1 hour at room temperature (RT). Slides were mounted using ProLong Gold Antifade reagent (ThermoFisher #P36934). Images were acquired at 20X magnification using the Zeiss Axio Observer 7 microscope. A control without primary antibody was included for each animal. NF-H staining intensity was quantified in Fiji by measuring the mean gray value within the largest possible nerve area. The NF-H staining area represents the percentage of the nerve area covered by NF-H positive signal.

### Protein extraction

Muscle and nerve samples were homogenized in their respective RIPA buffers: for muscle, the buffer contained 150 mM NaCl, 50 mM Tris (pH 8), 0.5% sodium deoxycholate, 1% NP-40, and 0.1% SDS; for nerve, it contained 2% SDS, 25 mM Tris (pH 8.0), 95 mM NaCl, 2 mM EDTA, and 0.5% sodium deoxycholate. Both buffers were supplemented with 1 mM PMSF, 1 mM sodium orthovanadate, 5 mM sodium fluoride, and 1X protease inhibitor cocktail. Tissue samples were processed using a Precellys® Evolution Touch tissue homogenizer (Bertin Technologies) with two 20-second cycles at 6000 rpm and cleared by centrifugation. Protein concentrations were determined using the DC Protein Assay Kit (#5000116, BioRad).

### Western blotting

Proteins were resolved on 10% polyacrylamide gels prepared in-house according to standard protocols. For muscle samples, 10 µg of protein in 12 µL were loaded per lane; for nerve samples, 5–12 µg of protein in 20 µL were used. Electrophoresis was performed at 130 V for approximately 1 hour. Proteins were transferred to nitrocellulose membranes using the Trans-Blot Turbo RTA Mini Nitrocellulose Transfer Kit (#170-4270, BioRad) for 5–10 minutes at 2.5 A. Membranes were stained with Ponceau S, then blocked for 1 hour in TBS containing 5% nonfat dry milk and 0.1% Tween-20 (TBST; #P2287-500ML, Merck). Membranes were incubated overnight at 4 °C with primary antibodies (Supplementary Table 1 Reagents) diluted in TBST with 5% milk. After washing, membranes were incubated for 1 hour at RT with the appropriate HRP-conjugated secondary antibody. Signal detection was performed using enhanced chemiluminescence reagents (#32209, ThermoFisher Scientific), and images were acquired on an Amersham Imager 600 (GE Healthcare Life Sciences). Band intensities were quantified using Fiji. Data were normalized to Ponceau staining and the mean value of control group.

All original, uncropped western blot images used for quantification, including those not shown in the figures and their corresponding loading controls, are available in supplementary material.

### Statistical analysis

All statistical analyses and graph generation were carried out using GraphPad Prism software (version 10.0.2). Since previous studies reported no sex-related differences in the *Dnm2*^K562E/+^ mouse model^20^, data from male and female mice were combined for most analyses, except for body mass, while ensuring balanced representation of both sexes across groups as much as possible (see Supplementary Table 2: Statistics). Data distribution was assessed using the Shapiro-Wilk test. For normally distributed data with equal variances, one-way ANOVA was performed followed by Tukey’s post hoc test. When variances were unequal, a Brown-Forsythe and Welch ANOVA was applied, followed by Dunnett’s T3 post hoc test. When data followed a log-normal distribution, a log transformation was performed prior to analysis. Non-normally distributed datasets were analyzed using the Kruskal-Wallis test with Dunn’s post hoc test. Two-way ANOVA was used when two factors were assessed (e.g., body mass × sex; force × stimulation frequency; axon distribution × diameter category; body mass or hanging time × age) followed by Tukey’s or Bonferroni’s multiple-comparison tests. All pairwise comparisons were conducted, but only statistically significant differences are shown on the graphs. A p-value < 0.05 was considered significant. Graphs display individual data points with mean ± SD. Detailed information regarding tests used and sample sizes is provided in Supplementary Table 2.

### Ethical approval

All animal procedures were conducted in compliance with French and European regulations and were approved by the Com’Eth IGBMC-ICS institutional ethics committee (Illkirch, France) under authorization number APAFIS #43721-202304241436921.

## Results

### Severe and irreversible motor defects in the *DNM2*-CMT mouse model

To evaluate the therapeutic potential of nerve-targeted DNM2 supplementation, we used the severe *Dnm2^K562E/^*^SC-^ mouse model. These mice carry systemic heterozygosity for the K562E mutation and lack WT *Dnm2* specifically in Schwann cells. Previous studies showed that expression of the K562E allele alone is sufficient to support correct radial sorting and initiation of myelination at postnatal day 5. However, it is not sufficient to maintain the myelinated state of Schwann cells, as by postnatal day 24 these mice display a marked reduction in myelinated axons and severe electrophysiological deficits^21^. Notably, only ultrastructural and electromyography (EMG) analyses were reported in these mice; we thus performed here a comprehensive phenotypic characterization.

As AAV9-mediated gene therapies have proven effective in both demyelinating and axonal forms of CMT^31,32^, and intrathecal AAV9 delivery under the Schwann cell-specific MPZ promoter efficiently targets the peripheral nervous system^33^, we delivered murine DNM2 under the control of the rat MPZ promoter intrathecally using AAV9 at a dose of 3.1 × 10¹¹ genome copies per mouse, administered at 4 weeks of age after symptom onset (Supplementary Fig. 1). Behavioral analyses were performed at 7 weeks, followed by functional tests and tissue sampling at 8 weeks (Fig. 1A).

**Fig 1.**
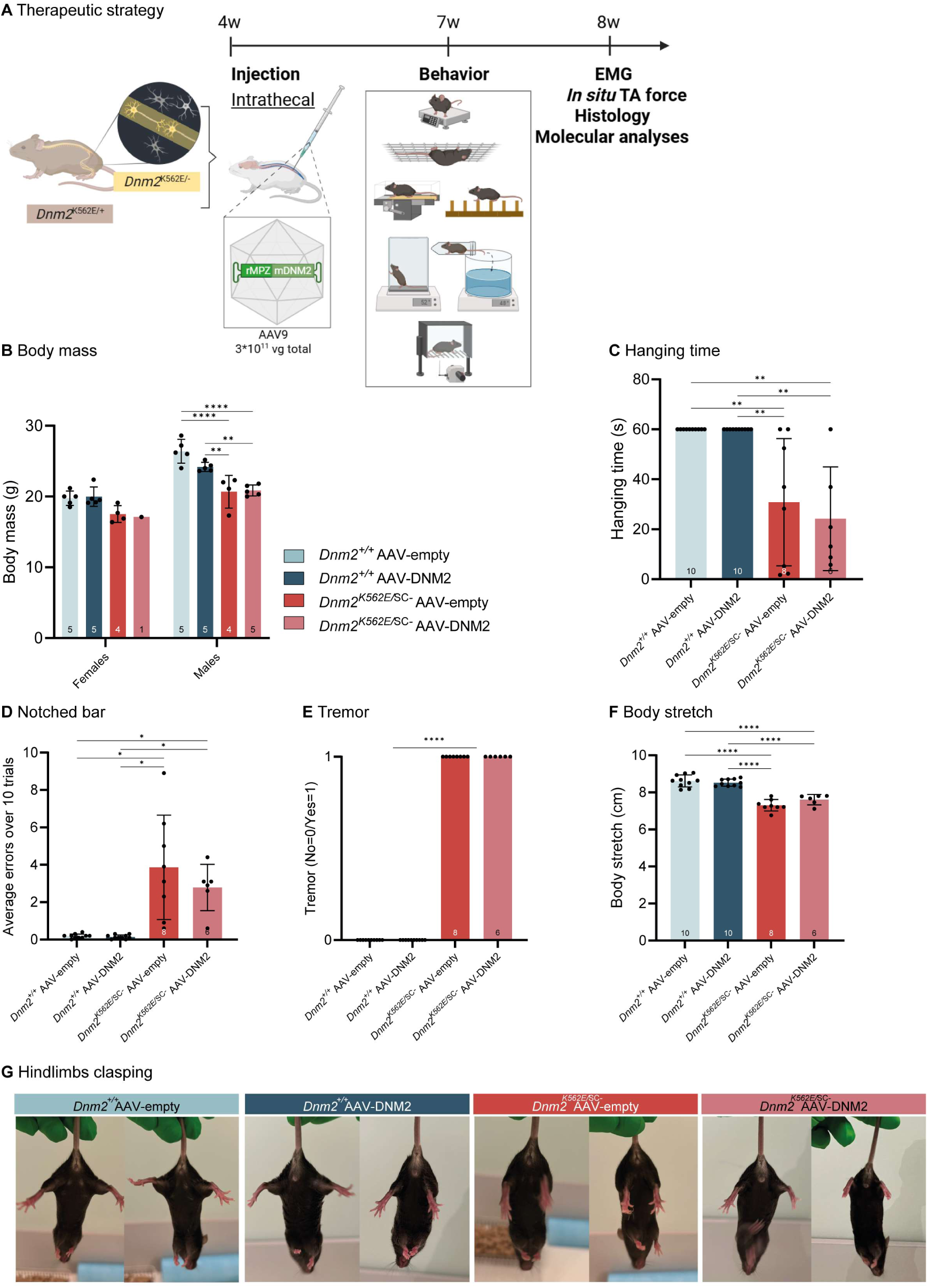
Severe and irreversible motor defects in the *DNM2*-CMT mouse model. **(A)** Experimental set-up. Created with BioRender. **(B)** Body mass of females (*Dnm2^+/+^*-empty, n=5; *Dnm2^+/+^*-DNM2, n= 5; *Dnm2^K562E/^*^SC-^-empty, n=4; *Dnm2^K562E/^*^SC-^-DNM2, n= 1) and males at 8w (*Dnm2^+/+^*-empty, n=5; *Dnm2^+/+^*-DNM2, n= 5; *Dnm2^K562E/^*^SC-^-empty, n=4; *Dnm2^K562E/^*^SC-^-DNM2, n= 5). Two-way ANOVA (interaction F=2.846 P=0.057, sex F= 69.13, P<0.0001, genotype F= 17.27 P<0.0001) with Tukey’s multiple comparisons. **(C)** Hanging test performance at 7w. Maximum hanging time = 60s. Kruskal-Wallis (P<0.0001) with Dunn’s multiple comparisons. **(D)** Average number of errors (hindlimb slips) when crossing the notched bar over 10 trials at 7w. Brown-Forsythe ANOVA test (F*=12.50, P=0.0011) with Dunnett’s T3 multiple comparisons. **(E)** Presence (1) or absence (0) of tremor at 7w. Chi-square test (Chi-square= 34.00, df=3, P<0.0001). **(F)** Body stretch (nose to tail base length) measured during treadmill walking at 7w. Ordinary one-way ANOVA (F= 47.24, P<0.0001) with Tukey’s multiple comparisons. (C-F) N: *Dnm2^+/+^*-empty, n=10; *Dnm2^+/+^*-DNM2, n= 10; *Dnm2^K562E/^*^SC-^-empty, n=8; *Dnm2^K562E/^*^SC-^-DNM2, n= 6. **(G)** Tail suspension at 7w, indicating presence or absence of hindlimb clasping (n=1 mouse per group). Each dot represents a mouse. Charts present individual values and mean ± standard deviation.*p<0.05, **p<0.01, ****p<0.0001.

First, we observed only a slight decrease in the mean DNM2 levels in the sciatic nerve of untreated *Dnm2^K562E/^*^SC*-*^ compared to controls (0.74-fold), likely due to the presence of other cell types (e.g., fibroblasts, endothelial cells, immune cells) (Supplementary Fig. 2A). Then, we validated by western blot a significant increase in DNM2 levels in the sciatic nerve of treated *Dnm2^K562E/^*^SC*-*^ mice (1.4-fold vs. controls, 1.9-fold vs. untreated *Dnm2^K562E/^*^SC*-*^). Unexpectedly, treated control WT mice showed a decrease in DNM2 expression, which does not preclude investigating the therapeutic potential of DNM2 overexpression in the *DNM2*-CMT model.

*Dnm2^K562E/^*^SC*-*^ mice exhibited reduced body size at 7 weeks (Supplementary Fig. 2B), and reduced body mass especially in males (Fig. 1B; Supplementary Fig. 1C). They also showed impaired motor performance, demonstrated by a twofold decrease in hanging time and increased coordination deficits, as indicated by a higher number of errors and falls on the notched bar test (Fig. 1C-D; Supplementary Fig. 1D and 2C). All *Dnm2^K562E/^*^SC*-*^ mice displayed tremor (Video 1, Fig. 1E). Despite these deficits, the *Dnm2^K562E/^*^SC*-*^ mice remained notably active in their cages and did not display signs of paralysis (Video 2). *Dnm2^K562E/^*^SC*-*^ mice showed a 1.3-cm reduction in body stretch during walking, consistent with shorter body length and increased stride length, further supporting coordination defects (Fig. 1F; Supplementary Fig. 2B and D). Hindlimb clasping during tail suspension was also observed, suggestive of underlying neurodegeneration (Fig. 1G). Thermal and mechanical sensitivity, assessed by hot plate, tail immersion, and Von Frey tests at 7 weeks, revealed no deficits in *Dnm2^K562E/^*^SC*-*^ mice (Supplementary Fig. 2E-G).

**Fig 2.**
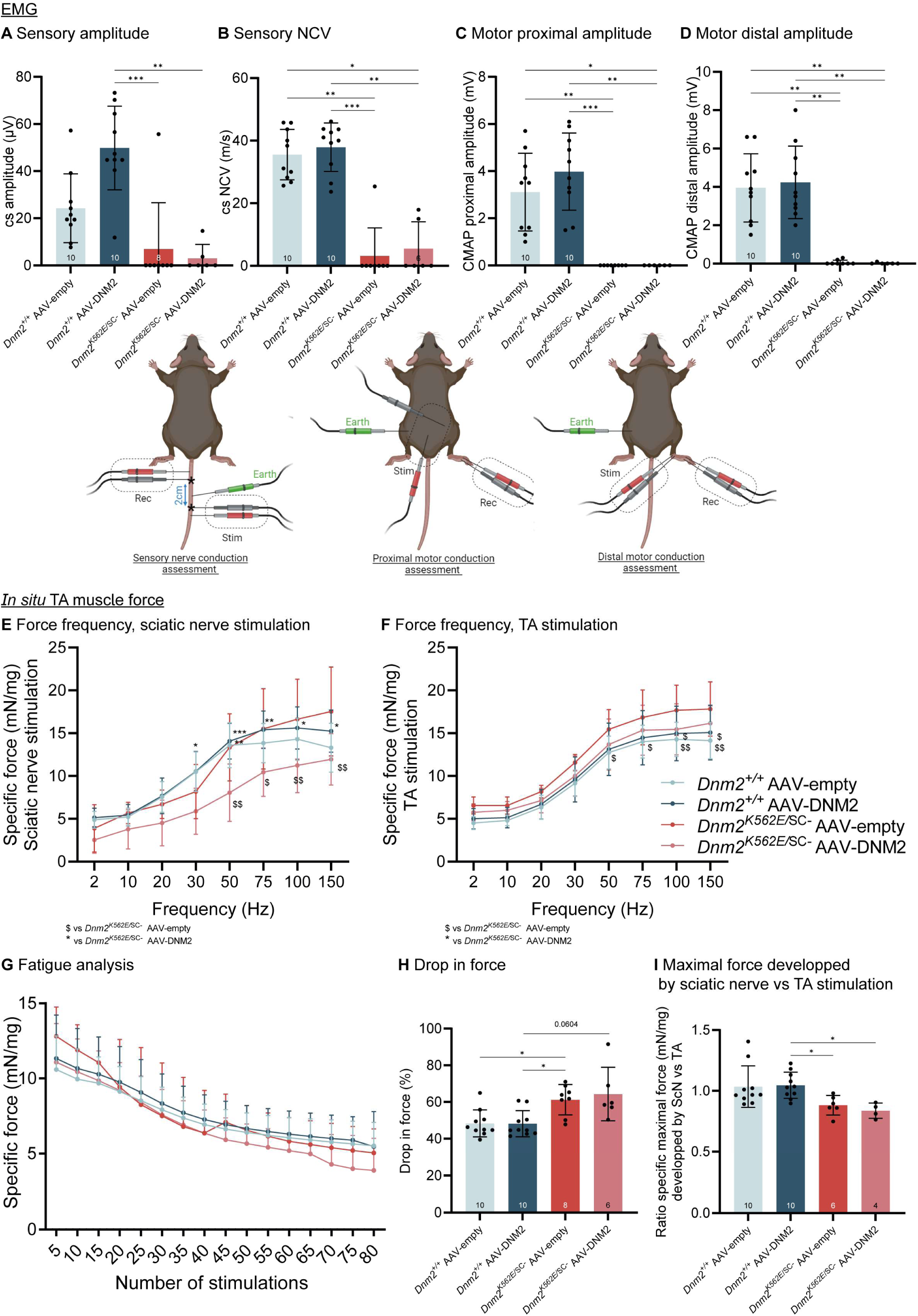
The *Dnm2^K562E/^*^SC*-*^ model exhibits severe CMT-like nerve conduction impairments. **(A-B)** Sensory electrophysiology at 8w: (**A**) compound sensory (cs) amplitude and, **(B)** nerve conduction velocity (csNCV). **(C-D)** Motor electrophysiology at 8w: **(C)** proximal compound muscle action potential (CMAP) and, **(D)** distal CMAP. Stim = stimulation, Rec = recording electrodes. Created with BioRender. (A-D) Kruskal-Wallis (P<0.0001) with Dunn’s multiple comparisons. **(E)** Force developed by TA muscle when stimulating the sciatic nerve. Two-way ANOVA (interaction F=1.152, P=0.2971, Frequency F= 75.15, P<0.0001, genotype F= 18.73, P<0.0001) with Tukey’s multiple comparisons. **(F)** Force developed by TA muscle when stimulating the TA at different incremental frequencies (1–150 Hz) at 8w, normalized to TA mass. Two-way ANOVA (interaction F=0.2002, P>0.9999, Frequency F= 142.7, P<0.0001, genotype F= 18.46, P<0.0001) with Tukey’s multiple comparisons. **(G)** Force developed by TA muscle over 80 consecutive TA stimulations at 40 Hz, normalized to TA mass. Two-way ANOVA (interaction F=0.4562, P=0.9991, Stimulations F= 38.20, P<0.0001, genotype F= 5.017, P=0.0020) with Tukey’s multiple comparisons. In (E-G), significant differences between *Dnm2^K562E/^*^SC-^ AAV-empty and other groups were indicated with $, while * shows differences between *Dnm2^K562E/^*^SC-^ AAV-DNM2 and other groups. **(H)** Drop in force between 5 first and 5 last stimulations during fatigue analysis at 40Hz. Kruskal-Wallis (P=0.0025) with Dunn’s multiple comparisons. **(I)** Ratio of the maximal force developed from sciatic nerve vs. TA muscle stimulation. Kruskal-Wallis (P=0.0035) with Dunn’s multiple comparisons. (A-D, F-H) N: *Dnm2^+/+^*-empty, n=10; *Dnm2^+/+^*-DNM2, n= 10; *Dnm2^K562E/^*^SC-^-empty, n=8; *Dnm2^K562E/^*^SC-^-DNM2, n= 6. (E, I) N: *Dnm2^+/+^*-empty, n=10; *Dnm2^+/+^*-DNM2, n= 10; *Dnm2^K562E/^*^SC-^-empty, n=6; *Dnm2^K562E/^*^SC-^-DNM2, n= 4. Each dot represents a mouse. Charts present individual values and mean ± standard deviation, *p<0.05, **p<0.01, ***p<0.001.

One month of DNM2 overexpression in peripheral nerves, initiated at 4 weeks of age, did not reverse the severe motor and coordination deficits in *Dnm2^K562E/^*^SC*-*^ mice, supporting that under these conditions of treatment these deficits are not reversible.

### The *Dnm2^K562E/^*^SC-^ model exhibits severe CMT-like nerve conduction impairments

To further investigate the neuromuscular dysfunction in this *DNM2*-CMT mouse model and evaluate potential rescue at the functional level following DNM2 supplementation, we next performed EMG and in situ muscle force measurements at 8 weeks.

EMG revealed markedly reduced or absent sensory nerve conduction amplitude and velocity in *Dnm2^K562E/^*^SC*-*^ mice (Fig. 2A-B). Proximal and distal motor responses were undetectable, preventing calculation of motor nerve conduction velocity (Fig. 2C-D). DNM2 increase in peripheral nerves through AAV-DNM2 intrathecal injection did not improve any of these electrophysiological parameters.

In situ force measurements revealed no relevant difference in Tibialis Anterior (TA) muscle force upon sciatic nerve stimulation among the groups (Fig. 2E). When stimulating the muscle directly, force output appeared slightly higher in mutants compared to controls, but this was likely due to normalization to a reduced muscle mass caused by atrophy (Fig. 2F). Fatigue exercise showed a greater decline in force production throughout the test in untreated mutants compared to controls (Fig. 2G-H). Although no intrinsic muscle weakness was evident, force generated through sciatic nerve stimulation tended to be lower than via direct muscle stimulation in *Dnm2^K562E/^*^SC*-*^ mice (Fig. 2I), suggesting conduction impairments.

Taken together, functional assessments showed no intrinsic muscle weakness but clear nerve conduction impairments in *Dnm2^K562E/^*^SC*-*^ mice, with no improvement following DNM2 intrathecal delivery. The *Dnm2^K562E/^*^SC*-*^ mice thus reproduce a severe peripheral neuropathy alike in *DNM2*-CMT patients.

### The *Dnm2^K562E/^*^SC^*^-^*model reproduces myelination deficits and axonal pathology characteristic of *DNM2*-CMT

To characterize the cellular basis for the nerve conduction impairments in the severe *DNM2*-CMT model, we investigated the structure and molecular biomarkers of the peripheral nerves. Semi-thin transversal sections of sciatic nerves stained with toluidine blue qualitatively revealed increased endoneurial connective tissue, signs of demyelination, and reduced fiber size (Fig. 3A). Quantifications showed a decreased number of total axons in the *Dnm2^K562E/^*^SC*-*^mice with no improvement upon DNM2 overexpression (Fig. 3B). Mutants sections exhibit increased number of fibers in a promyelinating stage, defined by Schwann cell nuclei positioned directly adjacent to axons (yellow arrow), a higher number of non-myelinated axons (purple arrow) almost absent in control groups, and a decrease number of myelinated axons (blue arrow). Among myelinated axons, their overall number was reduced in mutants, but the frequency of structural myelin abnormalities (orange and pink arrows) was normal. DNM2 supplementation did not improve these histological defects.

**Fig 3.**
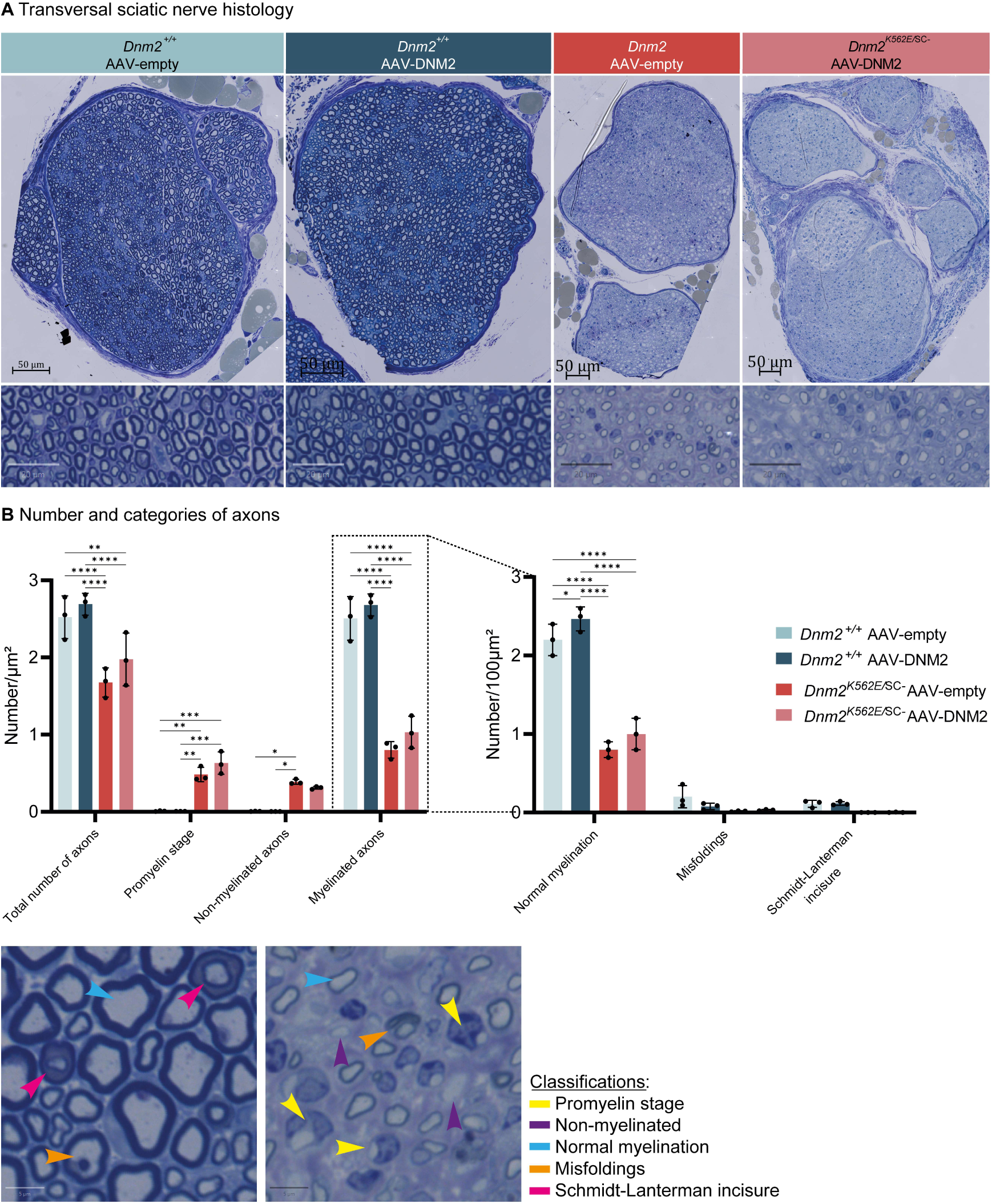
The *Dnm2^K562E/^*^SC*-*^ model reproduces myelination deficits and axonal pathology characteristic of *DNM2*-CMT. **(A)** Transversal section of the entire sciatic nerve stained with toluidine blue (upper panel, scale bar = 50 µm); magnified view in lower panel (scale bar = 20 µm). **(B)** Categories of axons in transversal sciatic nerve sections and their quantification per 100µm² at 8w. n=3 mouse per group. Two-way ANOVA (left graph: interaction F=40.26, P<0.0001, axon-type F= 468.8, P<0.0001, genotype F= 25.54, P<0.0001; right graph: interaction F=52.86, P<0.0001, axon-type F= 834.9, P<0.0001, genotype F= 81.15, P<0.0001) with Tukey’s multiple comparisons. Each dot represents a mouse. Charts present individual values and mean ± standard deviation, *p<0.05, **p<0.01, ***p<0.001, ****p<0.0001.

Among myelinated axons, the mean g-ratio 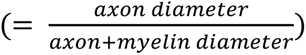 was increased in mutants, reflecting predominantly thinner myelin rather than reduced axon size, and this was not improved following DNM2 delivery (Fig. 4A-C). Scatter plots of g-ratio versus axon diameter showed consistently higher g-ratios in *Dnm2^K562E/^*^SC*-*^ mice across axon sizes compared to controls (Supplementary Fig. 3A), indicating relative myelin thinning. In controls, myelin thickness increased with axon diameter (∼17% per 1 μm), while in mutants this relationship was lost (slope near zero), suggesting disrupted coordination between axon size and myelin thickness (Supplementary Fig. 3B). In addition, the scatter plots revealed markedly fewer axons above ∼3-4 µm in *Dnm2^K562E/^*^SC*-*^ mice. Quantification confirmed a loss of large-caliber myelinated axons (Fig. 4D), similar to what is observed in patients. These parameters were unchanged following DNM2 supplementation.

**Fig 4.**
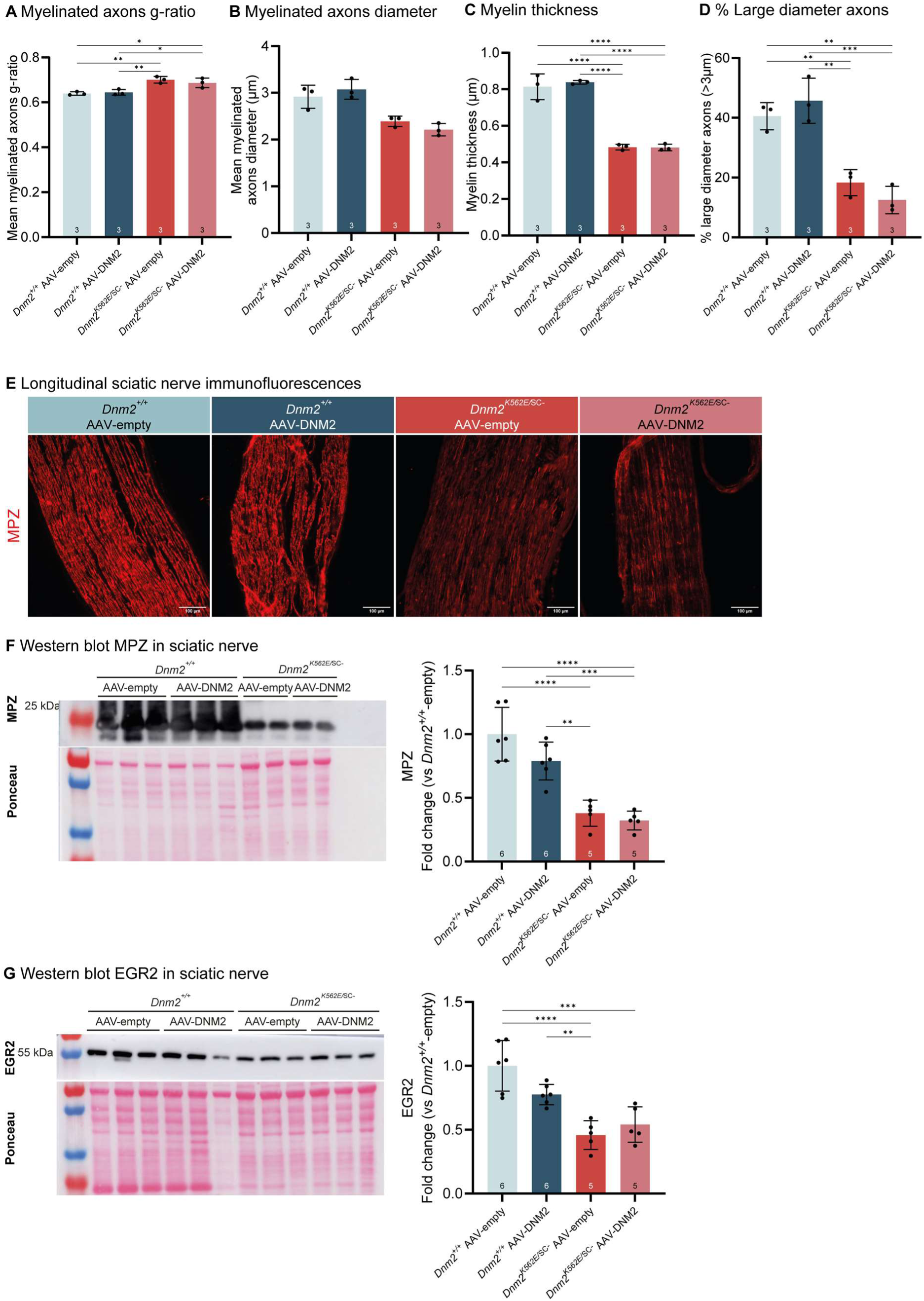
Defects in myelination are refractory to DNM2 supplementation. **(A)** Mean g-ratio 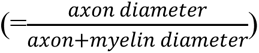 of myelinated sciatic nerve fibers at 8w. n=3 mouse per group. Ordinary one-way ANOVA (F= 12.56, P=0.0021) with Tukey’s multiple comparisons. **(B)** Mean axon diameter of myelinated sciatic nerve fibers at 8w. n=3 mouse per group. Kruskal-Wallis (P=0.0039) with Dunn’s multiple comparisons. **(C)** Mean myelin thickness of myelinated sciatic nerve fibers at 8w. n=3 mouse per group. Ordinary one-way ANOVA (F= 84.04, P<0.0001) with Tukey’s multiple comparisons. **(D)** Percentage of large axons (>3µm) in sciatic nerve sections. n=3 mouse per group. Ordinary one-way ANOVA (F= 27.11, =0.0002) with Tukey’s multiple comparisons. **(E)** Longitudinal sections of sciatic nerve immunolabeled for MPZ. Scale bar= 100µm. **(F)** Representative western blot and quantification of MPZ proteins level in sciatic nerves at 8w, normalized to Ponceau S staining. Ordinary one-way ANOVA (F= 26.49, P<0.0001) with Tukey’s multiple comparisons. **(G)** Representative western blot and quantification of EGR2 proteins level in sciatic nerves at 8w, normalized to Ponceau S staining. Ordinary one-way ANOVA (F= 16.73, P<0.0001) with Tukey’s multiple comparisons. (F-G) N: *Dnm2^+/+^*-empty, n=6; *Dnm2^+/+^*-DNM2, n= 6; *Dnm2^K562E/^*^SC-^-empty, n=5; *Dnm2^K562E/^*^SC-^-DNM2, n= 5. Each dot represents a mouse. Charts present individual values and mean ± standard deviation, *p<0.05, **p<0.01, ***p<0.001, ****p<0.0001.

Overall, the severe nerve conduction impairments of the *Dnm2^K562E/^*^SC*-*^ mouse is due to a decrease in myelination and axon loss.

### Molecular profiling reveals defects in myelination and inflammation that are refractory to DNM2 supplementation

To identify the molecular alterations underlying these structural defects, we performed several molecular analyses. Immunofluorescence on longitudinal sciatic nerve sections revealed reduced MPZ signal intensity in *Dnm2^K562E/^*^SC*-*^ mice compared to controls (Fig. 4E). This was confirmed by western blot, which showed a 62% decrease in MPZ protein levels compared to controls (Fig. 4F), consistent with the myelin thinning observed (Fig. 4C) and impaired neural transmission (Fig. 2A-D). We also detected a 54% reduction in the transcription factor EGR2 compared to controls (Fig. 4G). Given its role in activating myelin genes such as MPZ^34^ and in promoting the transition of promyelinating Schwann cells into fully myelinating cells^35^, its downregulation likely contributes to the demyelination and abnormal Schwann cell nuclear positioning observed. Neither MPZ nor EGR2 levels were restored by DNM2 overexpression.

We also performed immunofluorescence for NF-H (neurofilament heavy chain), a cytoskeletal component of large-caliber axons and a marker of axonal integrity (Fig. 5A, upper panel). Both staining intensity and NF-H-positive area tended to be reduced in treated and untreated *Dnm2^K562E/^*^SC*-*^ nerves compared to controls (Fig. 5B-C), consistent with the overall axon loss observed in transversal sections (Fig. 3B).

**Fig 5.**
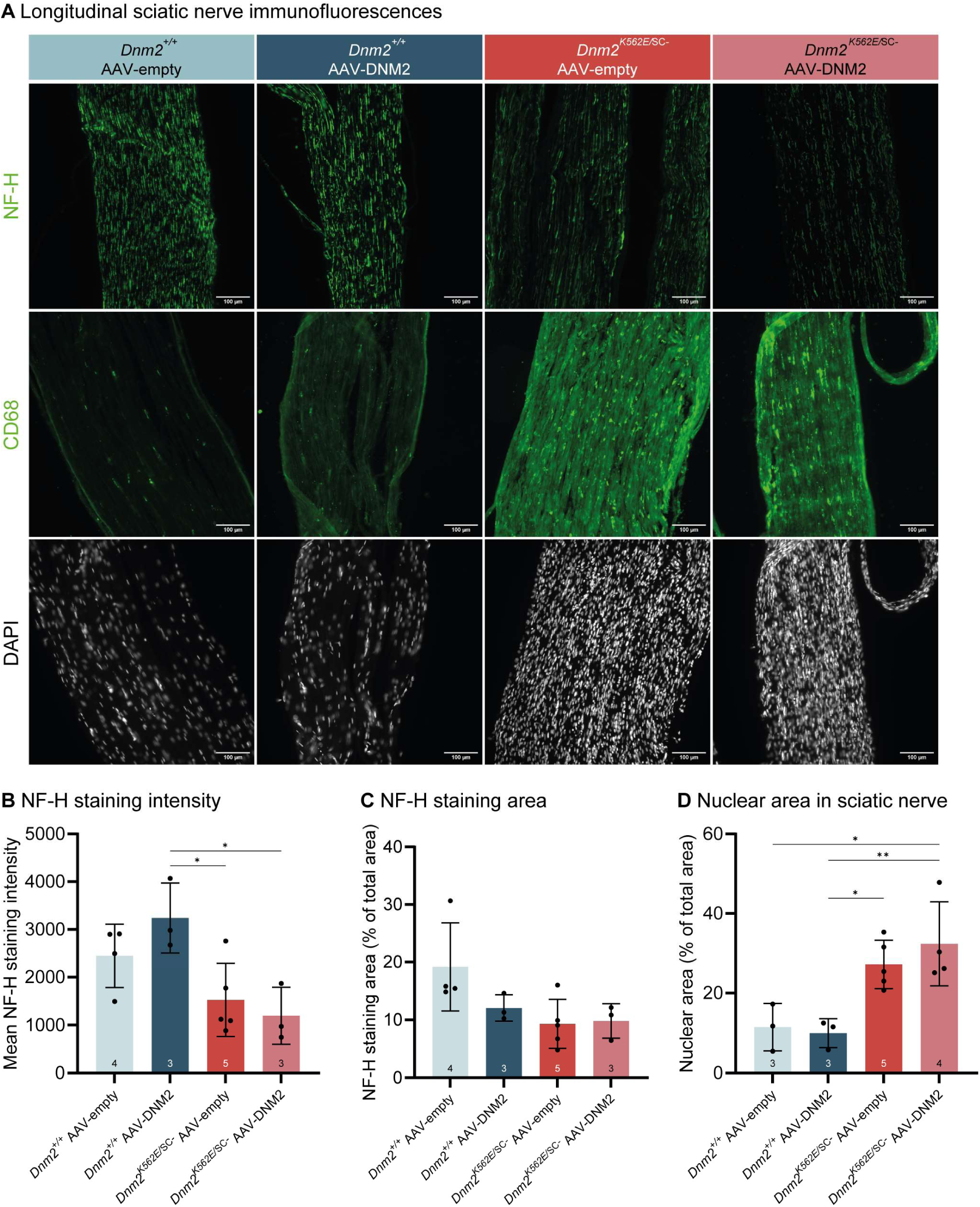
Schwann cell-specific DNM2 supplementation is insufficient to correct inflammation in a severe model of *DNM2-*CMT. **(A)** Longitudinal sections of sciatic nerves immunolabeled for NF-H (upper panel), CD68 (second panel), and DAPI (lower panel). Scale bar= 100µm. **(B-C)** Quantification of neurofilament-heavy (NF-H) staining **(B)** intensity (Ordinary one-way ANOVA (F= 5.702, P=0.0132) with Tukey’s multiple comparisons) and **(C)** area (Kruskal-Wallis (P=0.0618) with Dunn’s multiple comparisons) in longitudinal sciatic nerve sections. N: *Dnm2^+/+^*-empty, n=4; *Dnm2^+/+^*-DNM2, n= 3; *Dnm2^K562E/^*^SC-^-empty, n=5; *Dnm2^K562E/^*^SC-^-DNM2, n= 3. **(D)** Nuclear area in longitudinal sciatic nerve sections, calculated as DAPI-positive area relative to total nerve area. *Dnm2^+/+^*-empty, n=3; *Dnm2^+/+^*-DNM2, n= 3; *Dnm2^K562E/^*^SC-^-empty, n=5; *Dnm2^K562E/^*^SC-^-DNM2, n= 4. Ordinary one-way ANOVA (F= 8.432, P=0.0034) with Tukey’s multiple comparisons. Each dot represents a mouse. Charts present individual values and mean ± standard deviation, *p<0.05, **p<0.01.

To determine whether the observed nerve defects could correlate with inflammatory responses, we performed CD68 immunofluorescence, which revealed brighter signal intensity in *Dnm2^K562E/^*^SC*-*^ compared to controls (Fig. 5A, second panel). This was accompanied by a notable increase in nuclei density, indicative of immune cells infiltration (Fig. 5A, lower panel; Fig 5D). DNM2 intrathecal delivery did not reduce these inflammatory changes.

Together, these molecular findings highlight a combined defect in myelin, axonal maintenance, and inflammatory homeostasis in the peripheral nerves of *Dnm2^K562E/^*^SC*-*^ mice, that were not rescued by DNM2 overexpression.

### The *Dnm2^K562E/^*^SC^*^-^*model displays muscle atrophy that persist despite nerve-targeted DNM2 supplementation

Since myelination deficits impair neural transmission from motor neurons, which can lead to neuromuscular junction (NMJ) disruption and secondary muscle pathology, we evaluated both NMJs and muscle tissue. NMJs integrity was assessed using α-bungarotoxin labeling of Extensor digitorum longus (EDL) muscle (Fig. 6A). No signs of fragmentation were revealed, but a non-significant trend toward increased NMJ area was observed in mutant groups compared to controls (Fig. 6B), potentially reflecting compensatory remodeling. Muscle mass was reduced in both the TA and Gastrocnemius muscles of *Dnm2^K562E/^*^SC*-*^ mice (Fig. 6C-D). HE staining of TA sections revealed pronounced myofiber hypotrophy (Fig. 6E), confirmed by an increased proportion of small fibers (Fig. 6F-G). Western blot analysis showed no increase in DNM2 levels in the EDL muscle, which lies adjacent to the TA, confirming limited diffusion of the AAV from the intrathecal injection (Fig. 6H). None of the muscle phenotypes were improved by Schwann cell-specific DNM2 overexpression. It is thus possible that part of the motor phenotype observed in this severe *DNM2*-CMT model stems from primary muscle defects.

**Fig 6.**
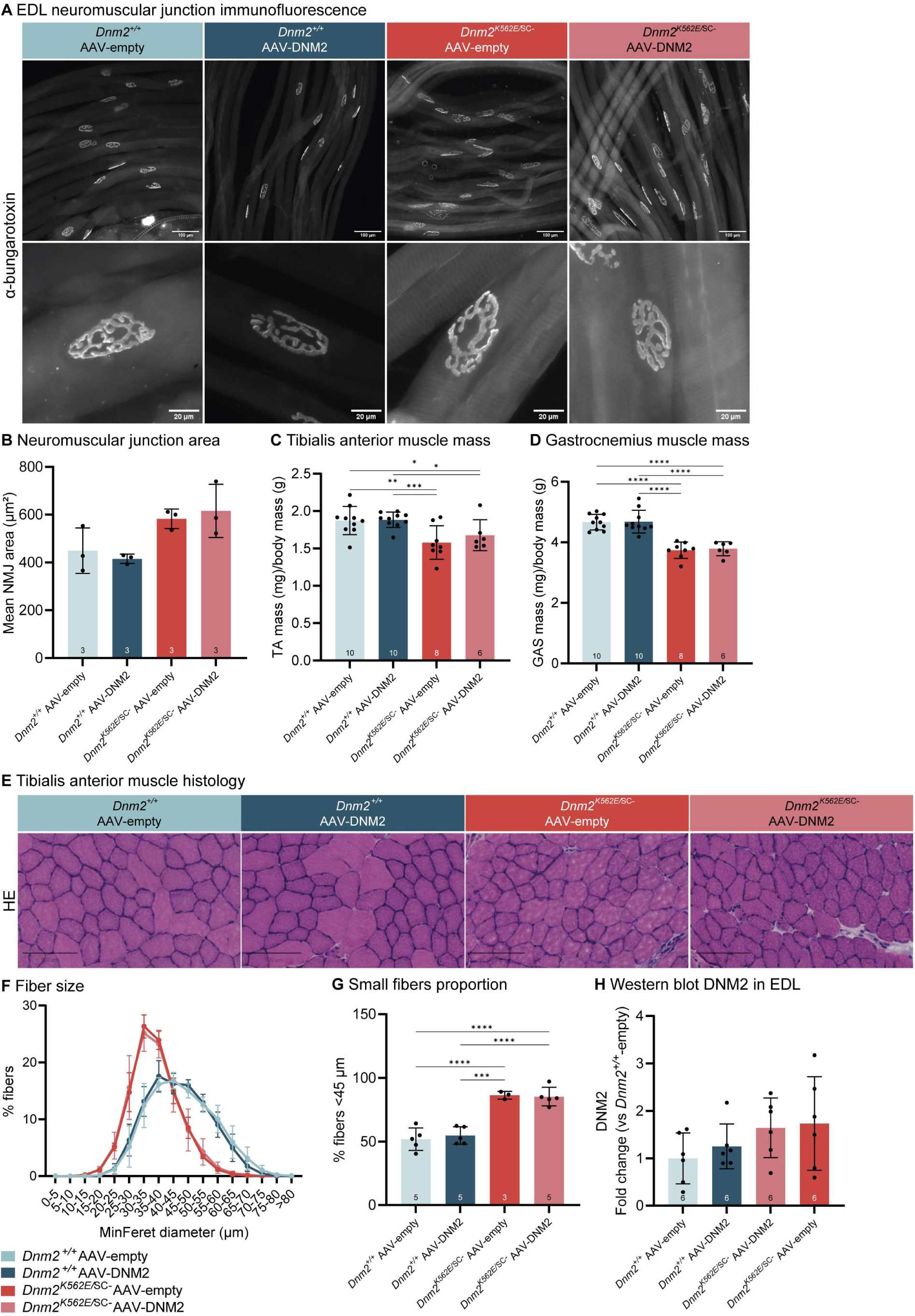
The *Dnm2^K562E/^*^SC*-*^ model displays muscle atrophy that persist despite nerve-targeted DNM2 supplementation. **(A)** Fluorescent α-bungarotoxin labeling of isolated extensor digitorum longus (EDL) muscle fiber bundles. Upper panel, scale bar = 100 µm; lower panel, scale bar = 20 µm. **(B)** Mean neuromuscular junction (NMJ) area in EDL muscle. n= 3 mouse per group. Ordinary one-way ANOVA (F= 4.928, P=0.0317) with Tukey’s multiple comparisons. **(C-D)** Muscle mass of **(C)** TA (Ordinary one-way ANOVA (F= 6.210, P=0.0021) with Tukey’s multiple comparisons), and **(D)** Gastrocnemius (Ordinary one-way ANOVA (F= 26.23, P<0.0001) with Tukey’s multiple comparisons) normalized to body mass at 8w. *Dnm2^+/+^*-empty, n=10; *Dnm2^+/+^*-DNM2, n= 10; *Dnm2^K562E/^*^SC-^-empty, n=8; *Dnm2^K562E/^*^SC-^-DNM2, n= 6. **(E)** TA transversal sections stained with hematoxylin-eosin (HE). **(F)** TA fibers distribution based on their MinFeret diameter. **(G)** Proportion of small fibers (MinFeret<45 µm) in TA sections. Ordinary one-way ANOVA (F= 29.96, P<0.0001) with Tukey’s multiple comparisons. In (F,G) N: *Dnm2^+/+^*-empty, n=5; *Dnm2^+/+^*-DNM2, n= 5; *Dnm2^K562E/^*^SC-^-empty, n=3; *Dnm2^K562E/^*^SC-^-DNM2, n= 5. **(H)** Quantification of DNM2 protein in EDL muscle, normalized to Ponceau S staining depicted in supplementary material. n = 6 mouse per group. Ordinary one-way ANOVA (F= 1.449, P=0.2585) with Tukey’s multiple comparisons. Each dot represents a mouse. Charts present individual values and mean ± standard deviation, *p<0.05, **p<0.01, ***p<0.001, ****p<0.0001.

## Discussion

This study investigated the neuromuscular defects underlying CMT neuropathy caused by dominant negative loss-of-function mutations in *DNM2*, and evaluated the therapeutic potential of increasing DNM2 expression.

Patients with *DNM2*-CMT experience distal motor and sensory deficits associated with markedly reduced nerve conduction velocity. Our detailed characterization of the *Dnm2^K562E/^*^SC-^ mouse validates this model as a robust tool for studying the peripheral nerve pathomechanism of *DNM2*-CMT (Supplementary Fig. 4). Although it is not genetically identical to patients, because *Dnm2* is selectively removed in Schwann cells in addition to the ubiquitous K562E mutation, its phenotype closely mimics the human condition and much more faithfully recapitulates neuropathic features than the *Dnm2^K562E/^*^+^ mouse, which displays only mild symptoms and a predominantly myopathic component^21^. The *Dnm2^K562E/^*^SC-^ model reveals a coherent cascade of pathological events that can explain patients’ phenotype. Reduced levels of the transcription factor EGR2 impair the activation of myelin genes, including MPZ, resulting in defective myelination characterized by decreased MPZ abundance and altered g-ratios. Impaired myelin disrupts axon-glia interactions and leads to a progressive loss of large-caliber myelinated axons, which are critical for rapid signal conduction. This axonal loss, combined with myelin thinning, produces the profound sensory and motor nerve conduction impairments observed in the model. Ultimately, these molecular and structural defects converge to produce the severe motor deficits characteristic of *DNM2*-CMT. These data also support an essential requirement for DNM2 during early postnatal Schwann cell maturation.

The rationale for supplementation with wild-type DNM2 in *DNM2*-CMT is a conceptually sound strategy. Indeed, CMT mutations in DNM2 are loss-of-function^16,17^. In vivo, complete loss of DNM2 in Schwann cells, whether constitutive or induced, leads to profound peripheral nerve defects^18^, and expression of the K562E mutant allele alone is insufficient to sustain peripheral nerve integrity^21^. Also, DNM2 reintroduction rescues the severe demyelination observed in Schwann cells lacking DNM2 in vitro^36^. However, our in vivo findings indicate that post-symptomatic intervention is insufficient to reverse established neuropathy (Supplementary Fig. 4). Intrathecal DNM2 delivery at 4 weeks, a stage when motor deficits and severe demyelination are already present, failed to improve motor behavior, nerve conduction, or histological outcomes, despite robust transgene expression. This lack of efficacy suggests that the therapeutic window for effective DNM2 augmentation in Schwann cells may occur earlier, likely before or during the onset of active myelination, which begins around postnatal day 5 in mice^37^. Consistent with this, previous work has shown that tamoxifen-induced DNM2 deletion in Schwann cells (*Dnm2*^SC-/SC-^) at 8 weeks causes demyelination that can be reversed by remyelination from non-recombined cells^18^. In contrast, *Dnm2^K562E/^*^SC-^ mice lack normal DNM2 activity from early developmental stages, resulting in cumulative and potentially irreversible damage to myelin and axons. Therefore, our results may reflect both a missed developmental window and the presence of irreversible structural degeneration by the time of treatment. Alternative explanations include suboptimal promoter strength, AAV serotype, or insufficient recovery period.

Overall, our results provide an in-depth characterization of a severe *DNM2*-CMT mouse model, that closely recapitulates the key clinical and pathological features of the human neuropathy. This work establishes the essential role of DNM2 in peripheral nerve integrity and myelination, and demonstrates that DNM2 supplementation after symptom onset is insufficient to reverse established pathology. The lack of therapeutic efficacy likely reflects a missed developmental window for effective intervention. These findings underscore the need for earlier or more sustained approaches, and highlight that therapeutic strategies for *DNM2*-CMT may need to prevent, rather than repair, Schwann cell dysfunction. Together, these findings offer valuable guidance for refining future gene therapy strategies targeting *DNM2*-related CMT.

## Data availability

All source data supporting the findings of this study are provided with the article.

## Acknowledgments

The authors would like to thank Ueli Suter and the Mouse Clinical Institute (ICS, Illkirch) for providing the *Dnm2^K562E/+^* mouse. We acknowledge the support of the scientific platforms at the Institut de Génétique et de Biologie Moléculaire et Cellulaire (IGBMC), particularly Nadia Messaddeqq and Chadia Nahy for the preparation of semi-thin nerve sections, Elise Lefebvre and Paola Rossolillo for virus production, and the histology platform for performing tissue staining. We acknowledge the IGBMC imaging center, member of the national infrastructure France-BioImaging supported by the French National Research Agency (ANR-10-INBS-04).

## Supplementary material

Supplementary material is available at *Brain Communications* online.

