## Supplementary material for "*DNM2*-CMT neuropathy stems from disrupted Schwann cell function and shows limited therapeutic reversibility"

### Supplementary Materials

Supplementary Fig. 1 to 3.

Supplementary Table 1. Reagents used. Antibodies used for immunofluorescences and western blots. Primers used for PCR and qRT-PCR.

Supplementary Table 2. Number of mice used per test and statistical analysis performed. For each test, number total of mice and sex-repartition as well as statistical analysis.

Video 1 : *Dnm2*$^{K562E/\mathrm{SC-}}$ male mouse exhibiting tremor at 8 weeks.

Video 2 : *Dnm2*$^{+/+}$ females (left cage) and *Dnm2*$^{K562E/\mathrm{SC-}}$ males (right cage) at 5 weeks, demonstrating that mutant mice remain active despite motor impairments.

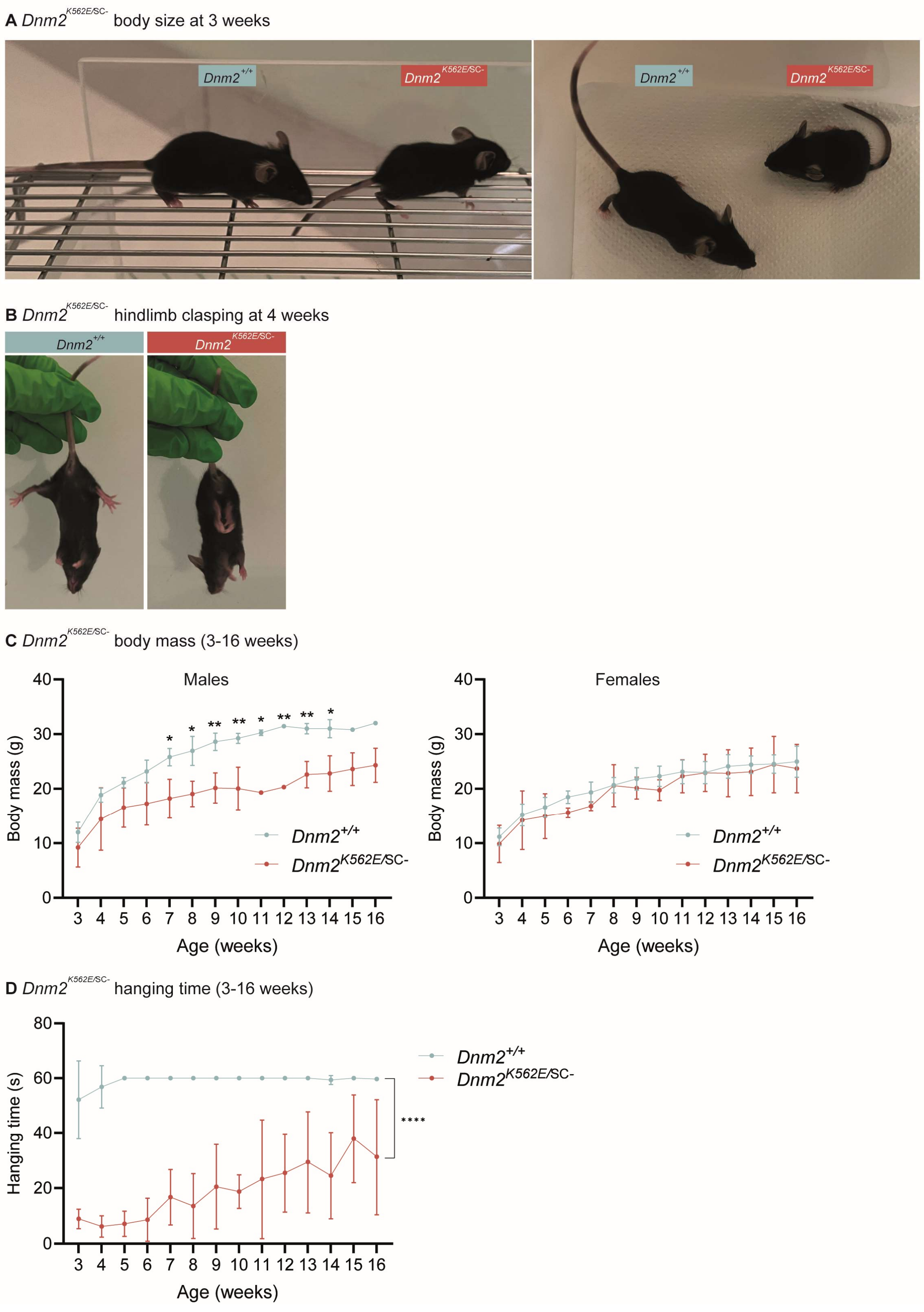


**Supplementary Fig 1. General characterization of the *Dnm2*$^{K562E/SC-}$ mouse line.** **(A)** Representative images of *Dnm2*$^{+/+}$ control and *Dnm2*$^{K562E/SC-}$ mice at 3 weeks. **(B)** Tail suspension at 4 weeks, indicating presence or absence of hindlimb clasping. **(C)** Body mass from 3 to 16 weeks. Males: *Dnm2*$^{+/+}$, n=3; *Dnm2*$^{K562E/SC-}$, n=2. Two-way ANOVA (interaction F=0.9447, P=5208, age F= 17.34, P<0.0001, genotype F= 127.6, P<0.0001) with Bonferroni's multiple comparisons. Females: *Dnm2*$^{+/+}$, n=3; *Dnm2*$^{K562E/SC-}$, n=4. Two-way ANOVA (interaction F=0.1337, P=0.9998, age F= 11.91, P<0.0001, genotype F= 4.007, P=0.0498) with Bonferroni's multiple comparisons. **(D)** Hanging time from 3 to 16 weeks. n=6 mouse per group. Two-way ANOVA (interaction F=2.228, P=0.0118, age F= 3.185, P=0.0004, genotype F= 594.8, P<0.0001) with Bonferroni's multiple comparisons. Charts present mean ± standard deviation, *p<0.05, **p<0.01, ****p<0.0001.

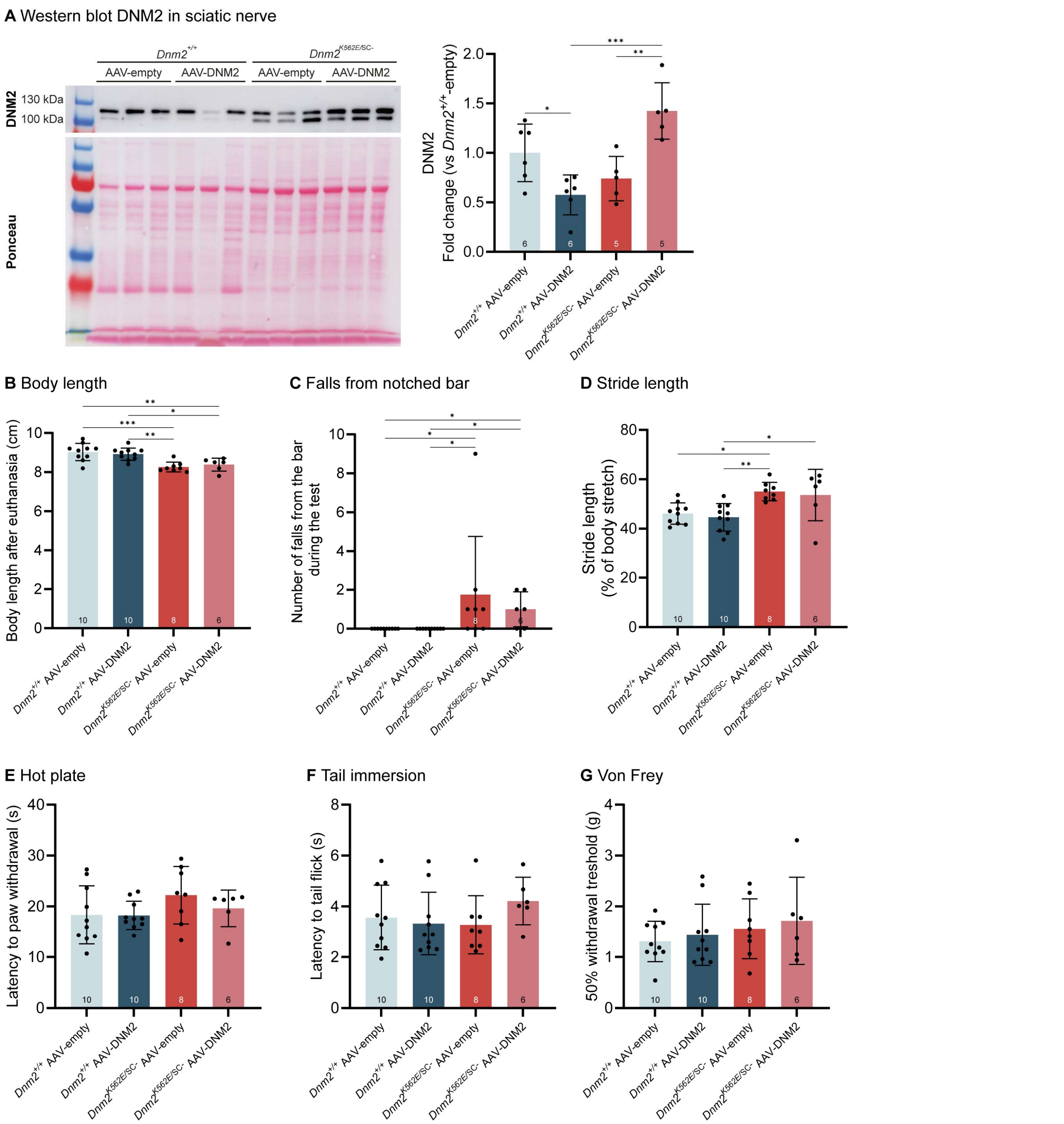


**Supplementary Fig 2. Validation of DNM2 overexpression in nerves and additional behavioral assessments. (A)** Representative western blot and quantification of DNM2 protein level in sciatic nerve at 8w, normalized to Ponceau S staining. *Dnm2*$^{+/+}$-empty, n=6; *Dnm2*$^{+/+}$-DNM2, n= 6; *Dnm2*$^{K562E/SC-}$-empty, n=5; *Dnm2*$^{K562E/SC-}$-DNM2, n= 5. Ordinary one-way ANOVA (F=11.33, P=0.0002) with Tukey's multiple comparisons. **(B)** Body length (nose to tail base) measured after euthanasia at 8w. Ordinary one-way ANOVA (F=10.28, P<0.0001) with Tukey's multiple comparisons. **(C)** Number of falls from the notched bar during the 10-trials test at 7w. Kruskal-Wallis (P=0.0009) with Dunn's multiple comparisons. **(D)** Stride (=length of a step) during treadmill walking normalized to body stretch at 7w. Ordinary one-way ANOVA (F=6.420, P=0.0017) with Tukey's multiple comparisons. **(E)** Hot plate test (52°C) at 7w. Reaction latency is recorded. Kruskal-Wallis (P=0.3408) with Dunn's multiple comparisons. **(F)** Tail immersion test (48°C) at 7w. Reaction latency is recorded. Ordinary one-way ANOVA (F=1.085, P=0.3706) with Tukey's multiple comparisons. **(G)** Von Frey test at 7w. Reaction to mechanic stimuli is recorded. The data presents the gram value at which the mouse is predicted to react 50% of the time. Ordinary one-way ANOVA (F=0.4963, P=0.6876) with Tukey's multiple comparisons. In (B-G) N: *Dnm2*$^{+/+}$-empty, n=10; *Dnm2*$^{+/+}$-DNM2, n= 10; *Dnm2*$^{K562E/SC-}$-empty, n=8; *Dnm2*$^{K562E/SC-}$-DNM2, n= 6. Each dot represents a mouse. Charts present individual values and mean ± standard deviation, *p<0.05, **p<0.01, ***p<0.001.

**A** Scatter plot of g-ratio as a function of axon diameter

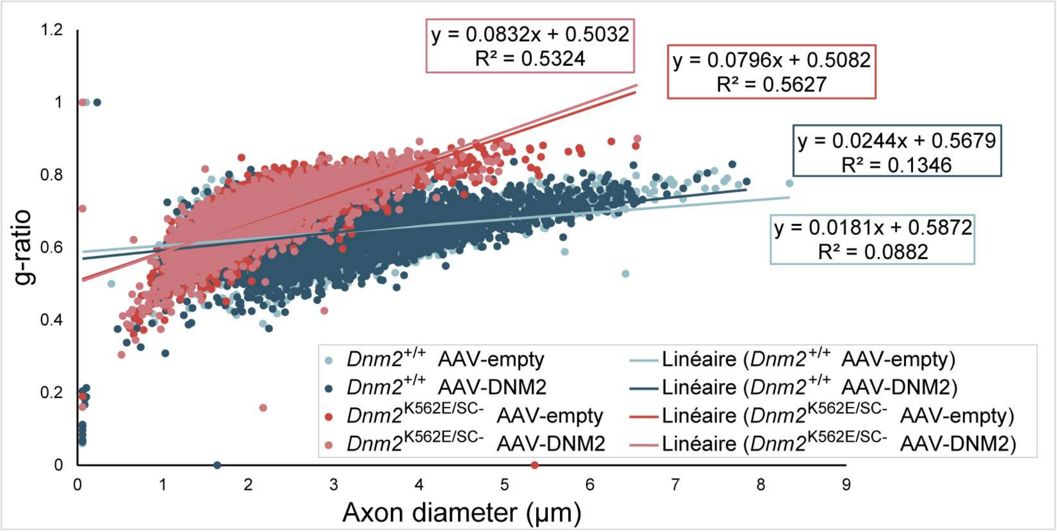


**B** Scatter plot of myelin thickness as a function of axon diameter

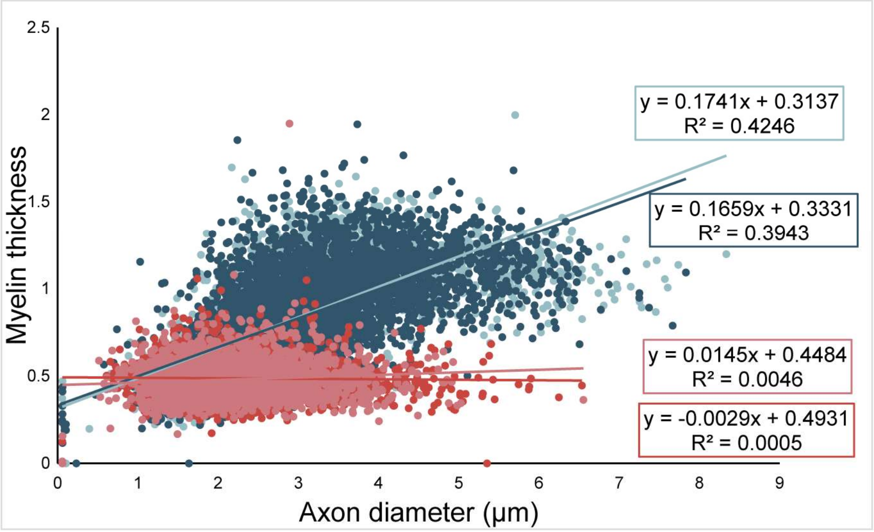


**Supplementary Fig 3. Relationship between axon diameter, g-ratio, and myelin thickness. (A-B)** Scatter plots of **(A)** g-ratio, and **(B)** myelin thickness as a function of axon diameter. *Dnm2*$^{+/+}$-empty, n=3 mice, 3447 fibers; *Dnm2*$^{+/+}$-DNM2, n= 3 mice, 4560 fibers; *Dnm2*$^{K562E/SC-}$-empty, n=3 mice, 2496 fibers; *Dnm2*$^{K562E/SC-}$-DNM2, n=3 mice, 2398 fibers; single fibers are plotted.

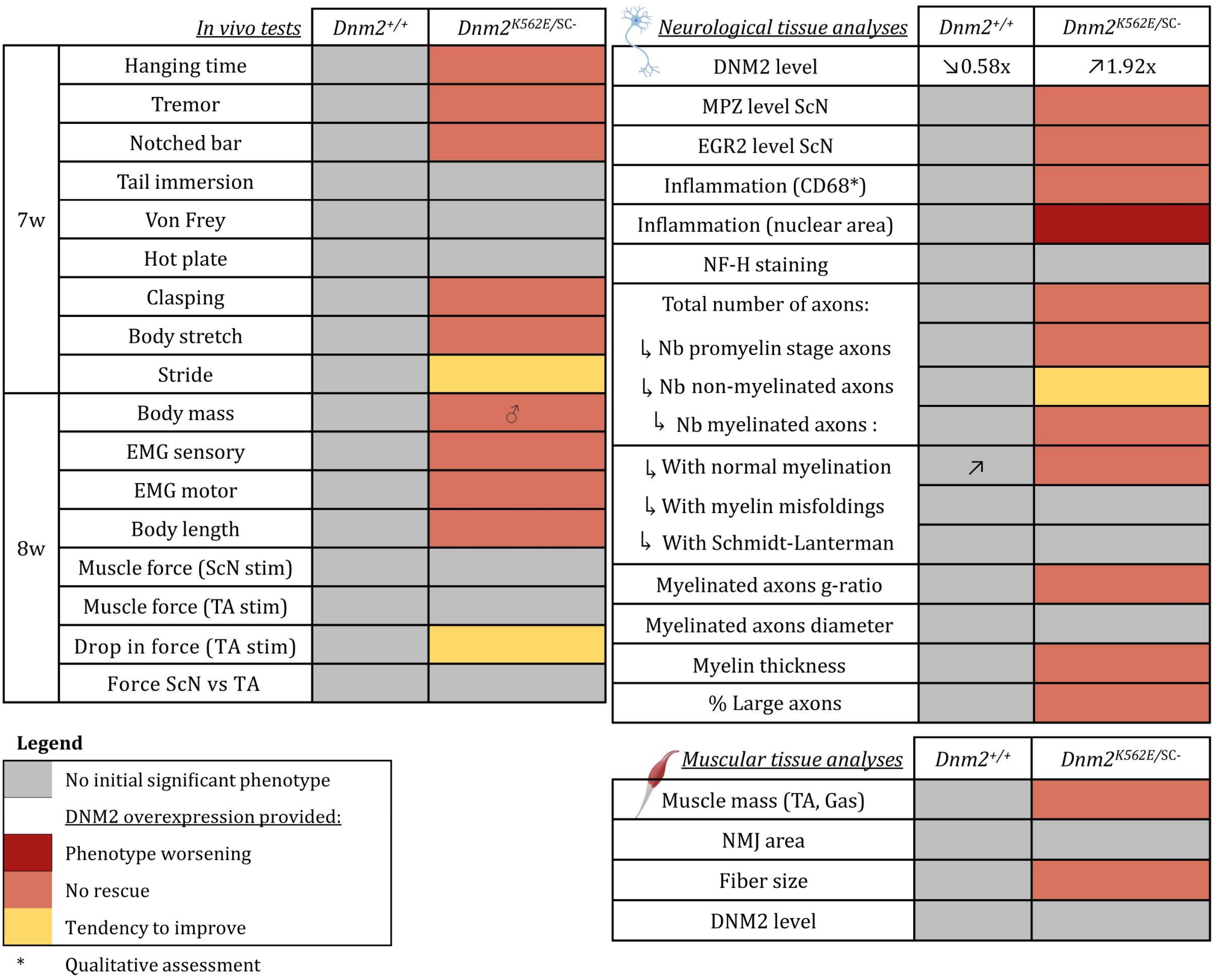


**Supplementary Fig 4. Overview of outcomes following DNM2 overexpression in *Dnm2$^{K562E/SC-}$* mice.** Summary of outcomes after intrathecal injection of murine DNM2 at 4w in *Dnm2$^{+/+}$* and *Dnm2$^{K562E/SC-}$* mice. Assessments were performed at 7-8w using in vivo tests and tissues analyses. TA= Tibialis anterior. ScN= sciatic nerve. Gas= Gastrocnemius. NMJ= neuromuscular junction. Tendency to improve= treated mutants are not different from both untreated mutants and controls.

| Reagent/Resource | Reference or Source | Identifier or Catalog Number |
|---|---|---|
| **Muscle immunofluorescence antibodies (dilution)** | | |
| CF488A Alpha-Bungarotoxin (1:1000) | INTERCHIM SA | 00005 |
| **Sciatic nerve immunofluorescence antibodies (dilution)** | | |
| MPZ, rabbit polyclonal (1:200) | Abcam | 31851 |
| ↳ GAR Alexa 555, goat polyclonal (1:250) | Invitrogen | A21430 |
| CD68, rat monoclonal (1:200) | Bio Rad | MCA 1957GA |
| ↳ GARat Alexa 488, goat polyclonal (1:250) | Thermo Scientific | A-11006 |
| DAPI (1:1000) | / | / |
| NFH, chicken polyclonal (1:500) | Abcam | 4680 |
| ↳ GAC Alexa 488, goat polyclonal (1:250) | Invitrogen | A11039 |
| **Western blot antibodies (dilution)** | | |
| DNM2, rabbit polyclonal (1:1000) | Homemade (2865) | N/A |
| MPZ, rabbit polyclonal (1:1000) | Abcam | 31851 |
| EGR2, rabbit monoclonal (1:1000) | Abcam | 108399 |
| GAR perox, goat polyclonal (1:10000) | Jackson Immunoresearch | 111-036-045 |
| **Genotyping PCR oligos** | 5'-sequence-3' | |
| 6115 Er KE | TACACTGTCTGCACTGTCGAGCCCTG | |
| 6116 Ef KE | GCCATCTTCAACACAGAGCAGAGGTG | |
| Cre 160 | GAACCTGATGGACATGTTCAGG | |
| Cre 161 | AGTGCGTTCGAACGCTAGAGCCTGT | |
| 4613 | GGCAGCAAGCCTGTTTACCCGC | |
| 4614 | GCTGAGCCTCACAGTGCAGAGCC | |
| **AAV titration qPCR oligos** | | |
| mDNM2 Fw | ACCCCACACTTGCAGAAAAC | |
| mDNM2 Rv | CGCTTCTCAAAGTCCACTCC | |
| CMVe For | TACGGTAAACTGCCCACTTG | |
| CMVe Rev | AGGAAAGTCCCATAAGGTCA | |

**Supplementary Table 1. Reagents used.** Antibodies used for immunofluorescences and western blots. Primers used for PCR and qRT-PCR.
